# Dynamic utilization of low-molecular-weight organic substrates across a microbial growth rate gradient

**DOI:** 10.1101/2021.12.01.470877

**Authors:** K. Taylor Cyle, Annaleise R. Klein, Ludmilla Aristilde, Carmen Enid Martínez

## Abstract

Constantly in flux, low-molecular-weight organic substances (LMWOSs) are at the nexus between microorganisms, plant roots, detritus, and the soil mineral matrix. Nominal oxidation state of carbon (NOSC) has been put forward as one way to parameterize microbial uptake rates of LMWOSs and efficiency of carbon incorporation into new biomass. In this study, we employed an ecophysiological approach to test these proposed relationships using targeted exometabolomics (^1^H-NMR, HR-LCMS) coupled with stable isotope (^13^C) probing. We assessed the role of compound class and oxidation state on uptake kinetics and substrate-specific carbon use efficiency (SUE) during the growth of three model soil microorganisms (*Penicillium spinulosum*, *Paraburkholderia solitsugae,* and *Ralstonia pickettii*) in media containing 34 common LMWOSs. Microbial isolates were chosen to span a gradient in growth rate (0.046-0.316 hr^−1^) and differ phylogenetically (a fungal isolate and two bacterial isolates). Clustered, co-utilization of LMWOSs occured for all three organisms, but temporal cluster separation was most apparent for *P. solitsugae*. Potential trends (p <0.05) for early utilization of more oxidized substrates were present for the two bacterial isolates (*P. solitsugae* and *R. pickettii*), but high variability (R^2^ > 0.15) and a small effect of NOSC indicate these are not useful relationships for prediction. The SUEs ranged from 0.16-0.99 and the hypothesized inverse relationship between NOSC and SUE was not observed. Thus, our results do not provide compelling support for NOSC as a predictive tool, implying that metabolic strategies of organisms may be more important than chemical identity in determining LMWOS cycling in soils.

**Importance:** Community-level observations from soils indicate that low-molecular-weight compounds of higher oxidation state tend to be depleted from soil solution faster and incorporated less efficiently into microbial biomass under oxic conditions. Here, we tested hypothetical relationships between substrate chemical characteristics and the order of substrate utilization by aerobic heterotrophs at the population-level in culture, using two bacterial isolates (*Ralstonia pickettii* and *Paraburkholderia solitsugae*) and one fungal isolate from soil (*Penicillium spinulosum*). We found weak relationships indicating earlier uptake of more oxidized substrates by the two bacterial isolates but no relationship for the fungal isolate. We found no relationship between substrate identity and substrate use efficiency. Our findings indicate that substrate chemical characteristics have limited utility for modeling the depletion of low-molecular-weight organics from soil solution and incorporation into biomass over broader phylogenetic gradients.

## Introduction

Low-molecular-weight organic substances (LMWOSs) represent the interface between the microbial cell and the decomposition of organic inputs into soil and have long been understood to be critical to soil processes (1, 2). This pool of soil organic carbon is a relatively small fraction of total dissolved organic carbon at any one point in time, with concentrations of individual compounds typically at less than 1 mM (3). Rapid cycling of LMWOSs involves their partitioning into microbial biomass, metabolic respiration to CO_2_, and contribution to the formation of mineral-associated organic matter (MAOM) (4, 5). The LMWOSs are released into solution through the action of exoenzymes depolymerizing particulate plant material and actively exuded by plant roots to shape soil microbial communities (6), thus altering soil solution nutrient composition (7). While there is an emerging understanding of the complex signaling roles of LMWOSs (8), they primarily serve as the currency of the subterranean economy, providing both the carbon and energy source to support heterotrophic microbial populations. Therefore, a mechanistic understanding of microbial uptake and transformation of LMWOSs is warranted as this is the process through which the majority of soil respiration is produced (9) and a pivotal step in the formation of MAOM (10).

In fact, microbial uptake of LMWOSs in soil solution is much faster than sorption onto soil particles, thus highlighting the importance of microbial processing on the long-term fate of LMWOS-derived carbon and nitrogen in soils (11). Uptake rates tend to be extremely rapid (< 30 min) and previous studies have indicated that the half-life of the compound in soil is influenced by the compound class (3, 12). More oxidized compounds, such as organic acids, have been shown to be removed from soil solution and appear in respired CO_2_ at faster rates than less oxidized compounds, such as sugars and amino acids (4, 13–16). Isotope (^13^C, ^14^C) tracer experiments have shown different affinity of cellular transport systems involved in uptake (17) by thousands of taxa (18).

Once transported within a microbial cell, LMWOSs are routed through metabolic pathways and incorporated into biomass production (anabolism, assimilation) or mineralized to CO_2_ while producing reducing equivalents (catabolism, dissimilation) (19, 20). Accurately modeling the partitioning between anabolism and catabolism, referred to as carbon use efficiency (CUE), is critical to forecasting future changes in the soil carbon sink (21–23). Carbon use efficiency has been observed to vary due to intrinsic physiological strategies (e.g., growth rate, genome size) as well as to external conditions (e.g., temperature, moisture, substrate quality, nutrient limitations) (24–26). At the scale of individual compounds, the internal energy content of the substrate being metabolized (denoted here as nominal oxidation state of carbon, NOSC) is one promising determinant of CUE (19, 27, 28). In this framework, due to energy limitations, the metabolism of the relatively more oxidized LMWOSs (higher NOSC) (e.g., organic acids) is hypothesized to result in lower assimilation efficiencies when these substrates are used as sole carbon sources. While the prevalence of oxygen in surface soils is typically considered to relieve thermodynamic limitations, recent field and laboratory observations have shown thermodynamics to still be a key regulator of aerobic respiration in carbon-limited aquatic systems (29–31) and this is supported by observations in intact soils as well (15).

Moreover, it is not likely that substrates are used sequentially by microbial populations when inhabiting environments containing diverse, low-concentration LMWOSs (32–34). Microbial metabolism may leverage the ability to route LMWOS-carbon from individual substrates divergently to anabolism or catabolism, thus producing specific substrate use efficiencies (SUE) that differ from relationships predicted from growth on a single substrate alone (19). Metabolomics using isotope labeling have observed nonuniform metabolic routing of assimilated substrates (35) and demonstrated the dynamic activation of specific metabolic pathways coinciding with the clustered, co-utilization of substrates in model organisms (36).

There is accumulated evidence that molecular size, compound class, and NOSC may be useful predictors uptake kinetics and use efficiency of LMWOS in soils and these concepts are being incorporated into substrate explicit modeling frameworks (37, 38). It is currently unclear whether correlations between compound characteristics and substrate utilization patterns (39) can be broadly applied to all aerobic heterotrophs in soil environments or whether these correlations simply arise from niche partitioning (e.g., faster-growing community members prefer to metabolize more oxidized substrates than slower-growing community members) (39). Therefore, there is a need to conduct ecophysiological studies to understand substrate utilization profiles and metabolic use across the breadth of microbial diversity in soil. Though reductionist by design, studies of this category are necessary for asking fundamental questions about the influence of substrate chemistry on the microbial metabolism of carbon and for providing parameter bounds to modelers. They may show utility for building predictive models of microbial community interactions (40) and complementing observations from the field when done under realistic conditions of substrate diversity and concentration (41).

Here the aim of our study was to investigate the role of compound class and NOSC on substrate utilization profiles, uptake rates, and use efficiency at the microbial population level. For this work, we have grown three microbial strains isolated from a forest soil in a defined media with 34 LMWOSs at realistic, equimolar concentrations (25 μM each). The chosen isolates represented distinct phylogenies, an ascomycete (*Penicillium spinulosum*) and two closely related to Betaproteobacteria (*Paraburkholderia solitsugae, and Ralstonia pickettii*), which exhibited a range of growth rates in the defined media. We put forth three hypotheses: (1) that there is a negative relationship between substrate NOSC and the midpoint of substrate uptake; (2) that there is a negative relationship between NOSC and SUE; (3) that individual SUEs would diverge widely from the cumulative CUE in accordance with co-utilization and segregated routing of carbon substrates within carbon metabolism. We employed time-resolved exometabolomics to characterize the depletion of compounds from extracellular solution during their growth. Here, substrate depletion in the extracellular media is assumed to be due to cellular uptake. Parallel experiments using treatments with selective isotopically labeled LMWOS were performed to determine SUE in a mixed-substrate media. Our findings provide insights on the utility of the inherent chemical characteristics of potential substrates as predictors of microbial preferences and usage efficiencies.

## Results

**Faster growth led to increased biomass production.** The chosen microbial isolates, which ranged in growth rate in minimal defined media, exhibited a normalized specific growth rate (μ) of 0.04-0.32 hr^−1^ (Table 1, Fig. S1). The bacterial isolates (*R. pickettii*, *P. solitsugae*) grew faster and exhibited much shorter lag times than the fungal isolate (*P. spinulosum*) (Table 1, Fig. S1). Despite having a longer lag phase than *P. solitsugae*, the fastest growing isolate, *R. pickettii*, produced the most biomass in the shortest time (Table 1). Accordingly, the biomass data collected from labeled growth trials show that *R. pickettii* produced more cellular biomass (in mg) per OD unit (higher *k*) than *P. solitsugae*. While *R. pickettii* and *P. spinulosum* had similar cellular carbon contents, *P. solitsugae* had substantially less carbon per cellular dry mass (Table 1). These results showed the choice of isolates represented a gradient in growth rates, with faster growth resulting in more biomass but not necessarily more biomass carbon.

**Table 1.**
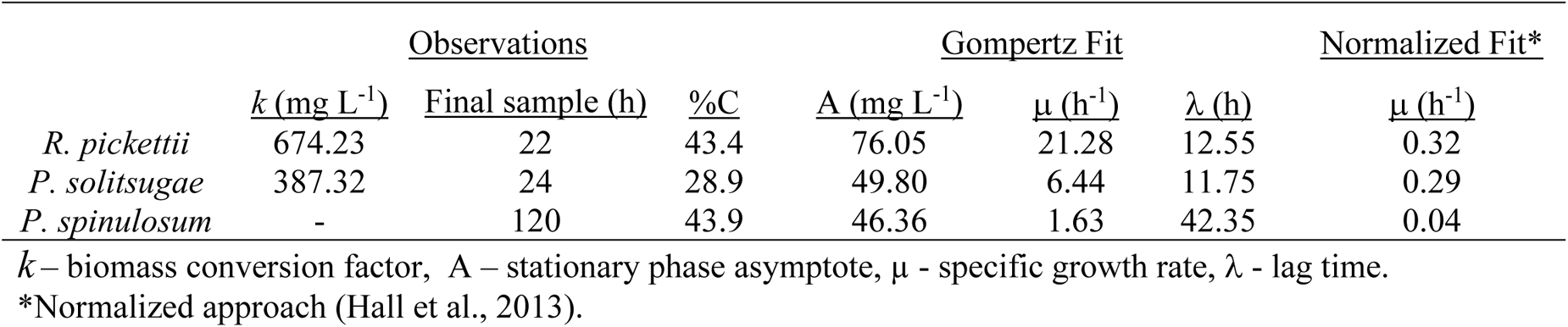
Observations and modeled growth curve fit parameters using biomass data.

|  | <u>Observations</u> |  |  | <u>Gompertz Fit</u> |  |  | <u>Normalized Fit*</u> |
| --- | --- | --- | --- | --- | --- | --- | --- |
|  | <u>k (mg L<sup>-1</sup>)</u> | <u>Final sample (h)</u> | <u>%C</u> | <u>A (mg L<sup>-1</sup>)</u> | <u>μ (h<sup>-1</sup>)</u> | <u>λ (h)</u> | <u>μ (h<sup>-1</sup>)</u> |
| *R. pickettii* | 674.23 | 22 | 43.4 | 76.05 | 21.28 | 12.55 | 0.32 |
| *P. solitsugae* | 387.32 | 24 | 28.9 | 49.80 | 6.44 | 11.75 | 0.29 |
| *P. spinulosum* | - | 120 | 43.9 | 46.36 | 1.63 | 42.35 | 0.04 |

*k* – biomass conversion factor, *A* – stationary phase asymptote, *μ* - specific growth rate, *λ* - lag time.
\*Normalized approach (Hall et al., 2013).

### Clustered LMWOS utilization in terms of compound class and NOSC

Throughout their growth, the three microbial isolates depleted nearly all LMWOS by at least 50% from the initial concentrations in the extracellular media (Fig. 1, Fig. S3-S5). The transposition of modeled substrate uptake inflection points of (*t_50_*) and usage time windows onto the growth curve of the isolate allowed comparisons of substrate usage patterns and usage overlap (Fig. 1). Using a k-means approach, three distinct uptake clusters (Cluster A, B, C) were identified for each isolate (Fig. 1A, 1C, 1E). The uptake of LMWOS occurred continuously whereby t_50_ values for substrates were distributed throughout the growth curve (Fig. 1B, 1D, 1F). Between the isolates, the clearest temporal separation of clusters was evident during substrate uptake by *P. solitsugae* (Fig. 1C), partially due to the presence of *t_50_* values early during the growth curve which were assigned to Cluster A (Fig. 1D). During the growth of *P. solitsugae*, a large proportion of each Cluster A substrate was depleted from the media (∼75%, 12 h) before significant depletion of any Cluster B and C substrates (Fig. 1D). Clustering of substrate uptake for *R. pickettii* (Fig. 1A) and *P. spinulosum* (Fig. 1E) produced uptake groups with less temporal distinction and *t_50_* values more closely centered around the midpoint of total carbon depletion than observed for *P. solitsugae* (Fig. 1D-E, Fig. S2). Greater than 70% of organic acid substrates were present in the earliest cluster for both bacterial isolates (Fig. 1B, 1D, Fig. 2A-B). Few compounds had significant early uptake by the fungus, *P. spinulosum,* with cluster A comprised of cysteine, gluconate, and the much slower uptake of glycine (Fig. 1F). In contrast to the bacterial isolates, *P. spinulosum* assimilated all organic acids except for gluconate in the last cluster, Cluster C (Fig. 1F, Fig. 2C). Clusters A and B were depleted completely for both bacterial isolates (Fig. 1A, 1C), while only Cluster B was depleted completely for *P. spinulosum* (Fig. 1E).

**FIG 1.**
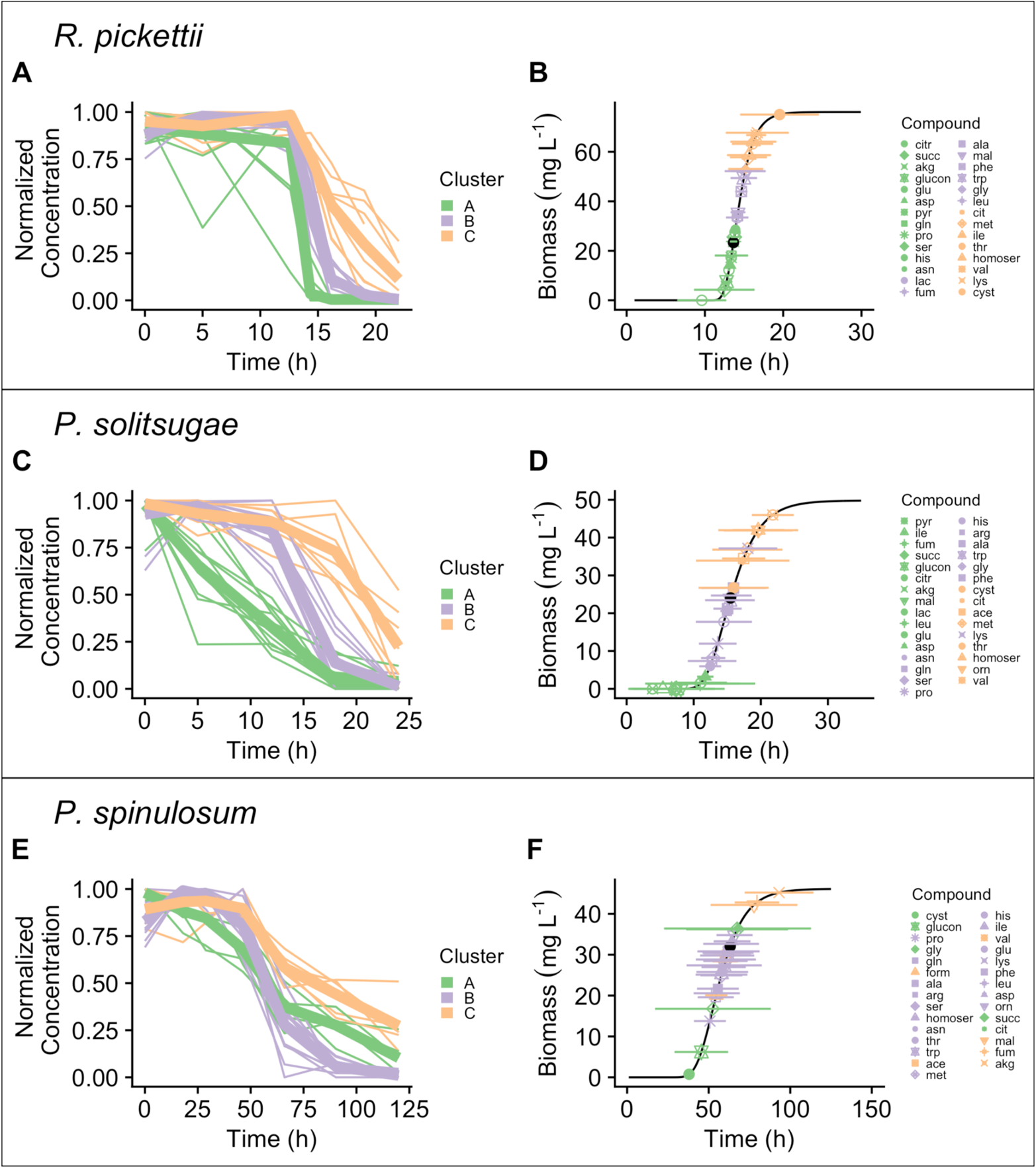
Depletion dynamics for observed LMWOS. Plots are paired by microbial isolate depicting depletion dynamics (A, C, E) and usage window plots (B, D, F) for uptake clusters identified using a k-means approach. Lines (A, C, E) are individual substrate uptake patterns normalized to initial concentration (mean, n = 3) with the thickest line showing the mean of the cluster. All points in usage window plots (B, D, F) are the mean modeled midpoints of depletion (*t_50_*) plotted over the growth curve of the isolate (black line). The mean midpoint of overall carbon depletion is shown with a black circle. Horizontal bars around each *t_50_* show the usage window (10% - 90% of initial concentration). All substrates are colored by cluster and listed in the legend in order of increasing *t_50_*. Ala, alanine; arg, arginine; asn, asparagine; cit, citrulline; cys, cysteine; glu, glutamate; gln, glutamine; gly, glycine; his, histidine; lys, lysine; met, methionine; orn, ornithine; ile, isoleucine; leu, leucine; lys, lysine; phe, phenylalanine; pro, proline; ser, serine; thr, threonine; trp, tryptophan; val, valine; ace, acetate; akg, α-ketoglutarate; form, formate; glucon, gluconate; lac, lactate; mal, malate; oxa, oxalate; pyr, pyruvate; succ, succinate.

**FIG 2.**
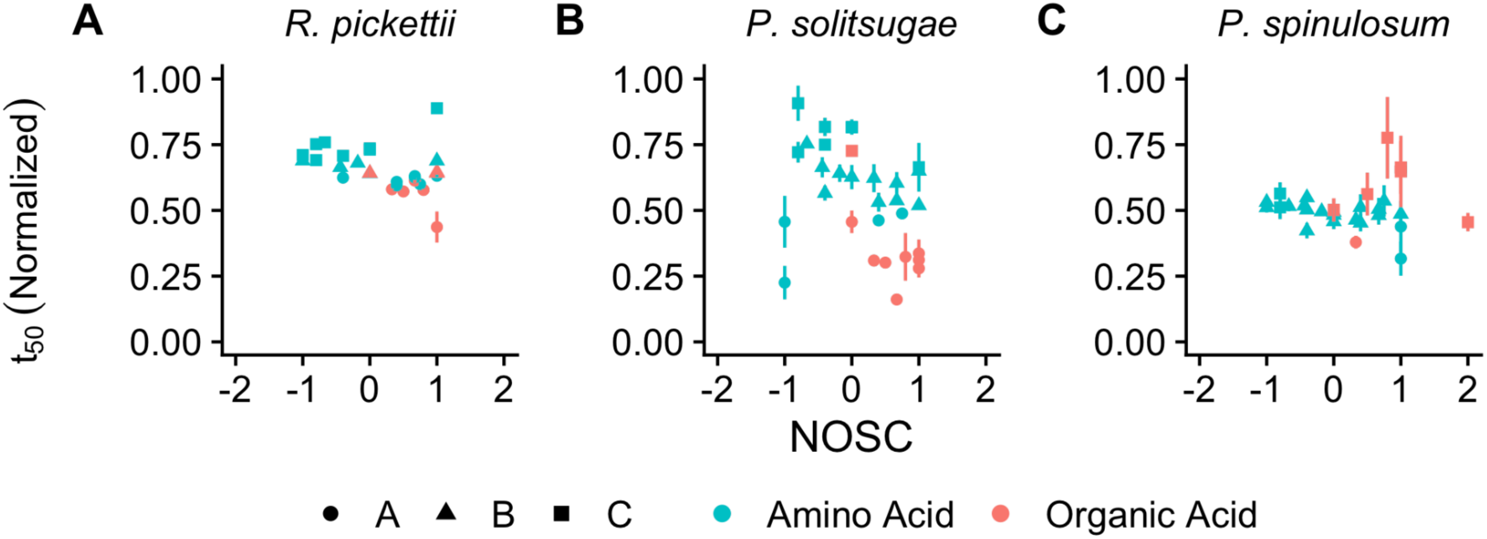
Inflection points of substrate depletion (*t_50_*) normalized to stationary phase sampling time as a function of substrate oxidation state (NOSC). Normalized *t_50_* values are the mean of three biological replicates with error bars representing standard error. Plots are paneled according to microbial isolate (A, B, C) and shapes illustrate cluster affiliation as determined using k-means clustering. Colors represent compound class (amino acid or organic acid).

We did not obtain compelling evidence of a correlation between substrate NOSC and substrate *t_50_* for any isolate (Fig. 2). The strongest potential trends were observed for *R. pickettii* and *P. solitsugae,* where the overall linear relationship between NOSC and the mean midpoint of substrate depletion (normalized *t_50_*) could be explained by the equations (normalized *t_50_* = 0.664 – 0.045 NOSC, adj R^2^ = 0.101, p = 0.05) and (normalized *t_50_* = 0.572 – 0.127 NOSC, adj R^2^ = 0.146, p = 0.019), respectively (Fig. 2A-B), These potential trends appear to be driven by the earlier average uptake of organic acids as compared to amino acids by both isolates (t-test results are p = 0.037, p < 0.001, respectively) (Fig. 2B-C). No significant correlation was observed for LMWOS uptake for *P. spinulosum* (Fig. 2C, p = 0.974), mostly due to the later uptake of three organic acids (malate, fumarate, α-ketoglutarate), though there was a trend for the earlier uptake of more oxidized amino acids (normalized *t_50_* = 0.489 – 0.040 NOSC, adj R^2^ = 0.232, p = 0.014). Despite a significant p-value (< 0.05) for some linear regression models in relation to the hypothesis of a correlation between NOSC and substrate utilization, all the linear regression models had a low adjusted R^2^ (< 0.25), which indicated high variability, and a small slope (< 0.13 NOSC), which indicated a relatively minor effect of NOSC unit on the midpoint of substrate depletion (regressions not shown).

Maximum substrate depletion rates decayed exponentially during population growth when displayed as a biomass normalized rate (Fig. 3A-C, Table S2-S4). Biomass-normalized substrate depletion rates were generally below 6.10 μmol h^−1^ mg_CDW_ for the two bacterial isolates (Fig. 3D-E, Table S2-S3), but the fungal biomass-normalized substrate depletion rates were much lower, predominantly below 0.15 μmol h^−1^ mg_CDW_ (Fig. 3F, Table S4). Though organic acids generally trended towards higher biomass-normalized depletion rates for the *P. solitsugae* (Fig. 3), no significant differences between depletion rates by compound classes were observed for any isolate.

**FIG 3.**
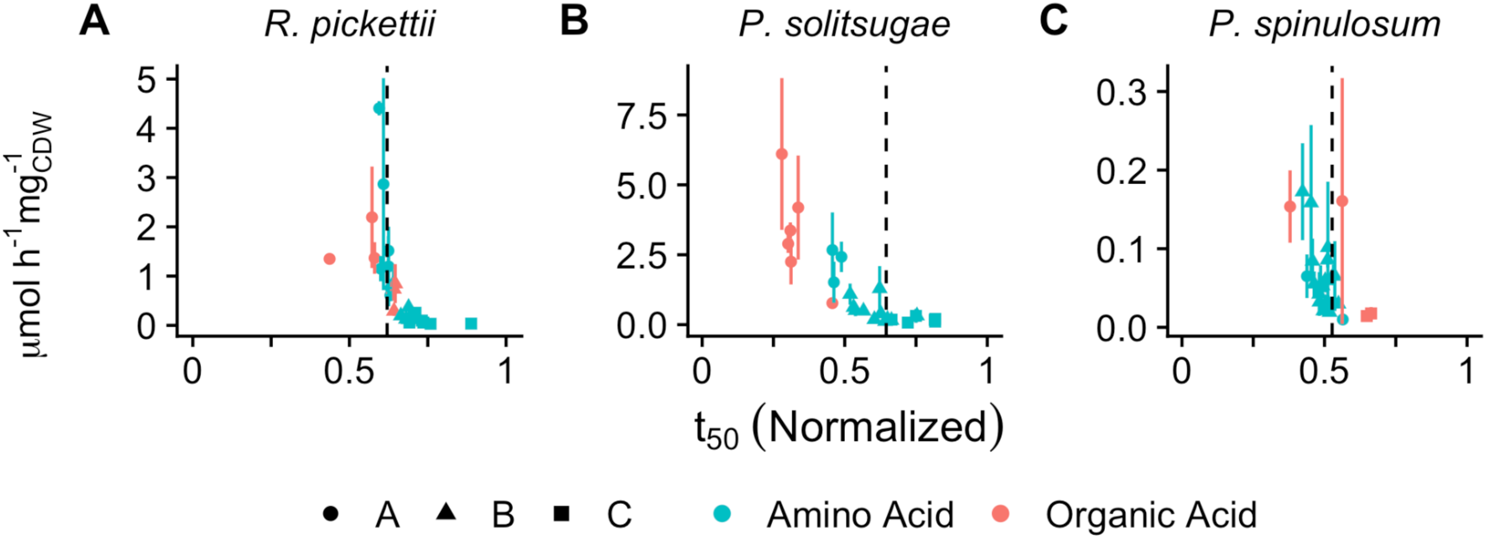
Biomass-normalized maximum depletion rate (A-C) as a function of the inflection point of depletion (*t_50_*) normalized to stationary phase sampling time. Values are the mean of three biological replicates with error bars representing standard error. Shapes illustrate cluster affiliation and colors represent compound class (amino acid or organic acid). The vertical dashed line depicts the normalized *t_50_* of overall carbon depletion from the media. Several points were removed for visualization purposes due to high variance between biological replicates (*P. solitsugae* – isoleucine, *P. spinulosum* – cystine & succinate).

### Substrate respiration dynamics, growth efficiency and substrate use efficiency of selected substrates

Labeled growth trials were used to track 5 specific substrates (glucose, acetate, formate, glycine, valine) that ranged in NOSC (−0.8 – 2) in the defined media (Fig. 4). All labeled substrates were respired (e.g., converted to CO_2_) to various degrees (Fig. 4A-C). The production of respired ^13^CO_2_ could be modeled in all cases except for *R. pickettii*’s respiration of ^13^C-formate (Fig. 4A, Table S5). Though insufficient data from early portions of the growth curve prevented fitting, the *t_50_* of ^13^C-formate respiration must have been less than 11.5 hours which was the first sampling time (Fig. 4A). In terms of initial media concentrations, the cumulative substrate-derived carbon that was respired from labeled substrates (CO_2_-C, *a*) straddled the proportion for all carbon sources for all isolates (Fig. 4A-C, Table S5). Cumulative respired CO_2_-C ranged from 12.74-25.29% of LMWOS-carbon present in the media (Fig. 4A-C, Table 5S). In all cases, carbon from the organic acids and sugar (acetate, formate, and glucose) appeared in respired CO_2_ earlier (lower *t_50_*) and more rapidly (smaller *w*) than the average of all carbon substrates (Fig. 4A-C, Table S5). The *t_50_* of respiration of these three substrates ranged from 0.19-6.07 h before the average of all substrates for bacterial isolates and 13.5-19.52 h for the fungal isolate (Fig. 4A-C, Table S5). The cumulative proportion of formate-carbon in respiration was at least 1.77-fold higher than any other substrate for all isolates and ranged from 19.84-59.19% of added substrate-carbon (Fig. 4A-C, Table S5). The two amino acids, glycine and valine, had respiratory curves that were later (higher *t_50_*) than overall CO_2_ production for all isolates (Fig. 4A-C, Table S5). This was most pronounced for valine for the two bacterial isolates, where valine was not respired until the beginning of stationary phase, representing a delay of greater than 3.09 h from the average of all LMWOSs respired, or greater than 16.3% of the total growth curve (Fig. 4A-B, Table S5). In contrast, respiration of glycine and valine much more closely followed overall respiration for *P. spinulosum*, with differences in *t_50_* ranging from 0.5-4.05 h or less than 4.2% of the total growth curve (Fig. 4C).

**FIG 4.**
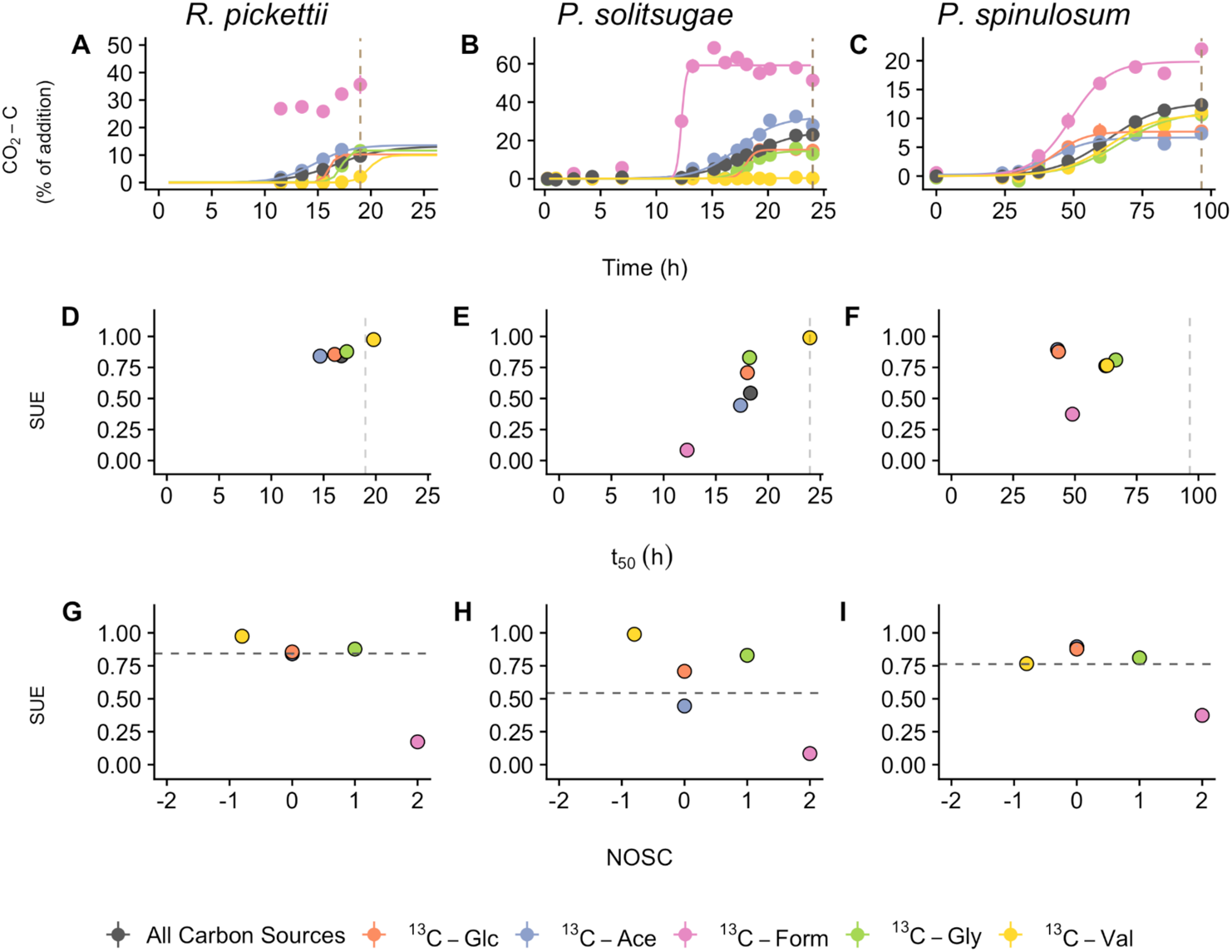
Respiration dynamics of ^13^C-labeled substrates and cumulative respiration along with estimates of substrate use efficiency (SUE). Graphs are grouped vertically for the three microbial isolates and arranged in order of decreasing specific growth rate (*R. pickettii > P. solitsugae > P. spinulosum*). Panels display respiration as a function of time in the first row of panels (A-C), SUE as a function of the midpoint of respiration, *t_50_,* (D-F), and SUE as a function of substrate oxidation state (G-I). The dashed vertical line in plots A-F indicates stationary phase of the overall growth curve and the sample point for determination of ^13^C present in microbial biomass. The dashed horizontal line in plots G-I indicates the cumulative carbon use efficiency of all carbon sources. See Fig. S6 for full panel A and Table S6 for microbial biomass carbon and nitrogen composition data. Glc, glucose; Ace, acetate; Form, formate; Gly, glycine; Val, valine.

Estimates of SUE produced values ranging from 0.16-0.99 (Fig. 4D-I, Fig. S7). The largest divergence in SUEs between highest and lowest efficiencies was found during the growth of *P. solitsugae* (Fig. 4E, 4H). For the two bacterial isolates, *R. pickettii* and *P. solitsugae*, substrates that appeared in respired CO_2_ earlier (lower *t_50_*) than the other substrates had relatively lower SUEs (SUE = 0.027 *t_50_* + 0.418, p = 0.025 and SUE = 0.078 *t_50_* - 0.807, p = 0.011, respectively) (Fig. 4D-E). There was no obvious trend between substrate respiration kinetics and SUE for *P. spinulosum* (Fig. 4F). Both *R. pickettii* and *P. spinulosum* exhibited a smaller range of SUE values, all of which (except formate) were at or near the overall CUE value, which was 0.84 and 0.76, respectively (Fig. 4G, 4I). These two isolates had a higher CUE than *P. solitsugae,* whose CUE was estimated at 0.54 (Fig. 4H). Formate had a significantly lower SUE for all isolates, never exceeding 0.31. No significant trends (p < 0.1) between the NOSC and SUE were found for any isolate (Fig. 4G-I).

## Discussion

This study aimed to characterize the substrate utilization profiles, uptake rates, and metabolic partitioning of carbon (CUE, SUE) of three soil microbial isolates with different growth rates. Using equimolar LMWOS concentrations and a diversity of substrates reflecting those present at the field site of isolation (32), we aimed to investigate the role of substrate energy content (NOSC) without confounding concentration differences. Stationary phase was chosen as the point of assessment for population-level carbon use efficiency (CUE_p_) to minimize the influence of secondary metabolite turnover on respiration measurements (21). While LMWOS are re-supplied in soil solution frequently, especially in the rhizosphere, these experiments mimic resource pulse events that induce microbial growth.

### Faster growth did not result in lower CUE

Specific bacterial phyla, such as Betaproteobacteria and Bacteroidetes, are known to grow rapidly in response to resource-pulses and this rapid growth is correlated positively to C-amendment levels in soil (42). Though the fungal community often represents a larger proportion of total microbial biomass (43), fungal:bacterial ratios derived from growth-based measurements made in C-units show bacterial dominance (F:B ranging from 0.02-0.44) in surface soils across a large gradient of ecosystem types (44). Rapid incorporation of labeled substrate pulses into the phospholipids of Betaproteobacteria also support the view that r-strategist fractions of the community are competitive LMWOS incorporators (39). In accordance with this paradigm, our growth data illustrated faster growth (Table 1) and higher substrate depletion rates (Fig. 3) by the two Betaproteobacteria compared to the ascomycete. Our growth rates in defined media were within the range of reported specific growth rates from intact soils (45–47). *P. solitsugae* grew faster in this defined media than in soil extract from the field site of origin (0.29 vs 0.17 hr^−1^) (32), indicating that estimated growth rates for these isolates are likely to be lower in intact soils.

Lowered rates in more chemically complex soil solution are potentially due to induced stress from other compounds present (e.g., antibiotics) or reduced bioavailability of LMWOS. Evaluations of fungal-to-bacterial dominance in long-term experiments (48) and laboratory assessments (44) have shown increased fungal response to high quality plant litter inputs, complicating the broad delineation that r-strategists (such as Betaproteobacteria) are the first responders to LMWOS inputs. Fungal species, such as *P. spinulosum*, may have competitive advantages in a complex soil environment other than growth rate, including the ability to explore more soil pore volume for available substrates (49). Faster growth is often assumed to come at the expense of CUE (24), representing a tradeoff between the ability to rapidly compete for available substrates and the efficiency of their incorporation into new cellular material.

Our estimates of cumulative CUE (Fig. 4G-I) did not align with theories of lower CUE from faster growing bacterial phylogenies (26, 50). The CUE estimates (0.54-0.84), calculated using biomass and CO_2_ datasets collected with population cultures of each organism, were about the same as or higher than the average estimate from soil microbial communities of ∼0.55 (24) and higher than CUE measurements taken at larger temporal or spatial scales, which range from ∼0.2-0.5 (51, 52). Though higher values are expected for measurements at the population level (CUE_p_) (21), our observations approached the theoretical maximum efficiency (0.88) of reduced compounds (19, 53). Lower CUEs in soil may result from increased biomass maintenance costs, nutrient limitation, and resource limitation (24, 54, 55), though certain discrepancies may be a result of methodological considerations (56). The defined media used in this experiment was not nutrient limited and all carbon substrates were in monomer form, likely providing favorable conditions for organisms to grow at their maximum potential efficiency. The correlation between growth rate and CUE has been shown to be complex and sometimes inconsistent (50). It was not therefore surprising that we did not observe any correlation between growth rate and CUE under these growth conditions. Furthermore, as described in a revised trait-based theory of microbial life history (Y-A-S), high growth rate may result from any ecological strategy paired with the right context and does not necessarily entail a tradeoff with yield (57).

### LMWOS utilization: Evidence for substrate co-utilization

Microbial substrate utilization preferences are thought to arise from the competitive advantage substrates offer to whichever ecological strategy is employed by the organism. Termed carbon catabolite repression, most microbes sequentially move through LMWOS that offer higher potential growth rates or yields (58, 59). Heterotrophic microorganisms have been known to deviate from sequential, diauxic growth when diverse substrates are present at low concentrations (34) and new exometabolomic work has stressed substrate co-utilization as a typical phenomenon (36, 40). Here, our data also provided evidence for diauxie and co-utilization in our substrate uptake dynamics for all isolates, characterized by clustered substrate uptake patterns with overlapping usage windows (Fig. 1).

The growth of the bacterial isolate, *P. solitsugae*, provided the strongest case for clustered substrate co-utilization, with broader substrate usage across the growth curve and more distinct cluster dynamics than the other two isolates (Fig. 1C-D). By comparison, *R. pickettii* and *P. spinulosum*, had more tightly grouped substrate clusters centered around the midpoint of total carbon depletion from the media (Fig. 1A-B, Fig. 1E-F, Fig. 3D, 3F). High resolution sampling along the growth curve may be needed to observe lags in growth, which would be indicative of changes in membrane transport system regulation as the population switched to a new cluster of substrates, such as in the case of *P. solitsugae*. Thus far, we have assumed that growth of these microbial populations relied on a relatively homogeneous metabolism within the population at any point in time. However, there could be anabolic heterogeneity within the population, as has been shown with single-cell studies (60), and this fluctuating heterogeneity could explain some of our divergence in substrate uptake between isolates. Phenotypic heterogeneity in terms of metabolism has been observed in clonal *Saccharomyces cerevisiae* populations under nutrient-limited conditions and is likely common in microbial populations (61, 62). Regardless of the mechanism leading to both diauxic and co-utilization patterns during substrate uptake, prior work with a marine heterotroph, *Pseudoalteromonas haloplanktis*, as well as two common soil pseudomonads indicate that potential substrate-specific growth rate may still dictate substrate preference (36, 40).

The ordering of LMWOS utilization could arise from the interplay between the expression of membrane transporter systems, system affinity, and the metabolic processing rates of the substrate inside the cell (17). We found substantial uptake of organic acids predominantly in the initial cluster (Cluster A) for both bacteria, but not for the fungal isolate (Fig. 2). These data are consistent with prior observations of the earlier uptake of organic acids compared to sugars and amino acids from soil solution by intact soil communities (4, 15), implying that this phenomenon is driven by fast-responding, bacterial populations similar to the two Betaproteobacteria examined in this study. Early organic acid uptake at the population-level in these bacteria may be due to the need for reducing equivalents from the metabolism of citric acid cycle intermediates when initiating growth, albeit not all organic acids (e.g., gluconate, formate, pyruvate) feed into the TCA cycle directly. The LMWOS substrate uptake ordering may also be dictated by transporter system expression more than the affinity of the different transporter systems. Our uptake rates (Fig. 3A-B) were comparable with the lowest rates measured for other soil bacterial species (40) and 2-3 orders of magnitude below modeled maximum substrate uptake rates (<74.9 µM substrate (mg C biomass)^−1^ hr^−1^ vs. 33,000 µM substrate (mg C biomass)^−1^ hr^−1^) (63). Fungal uptake rates were substantially lower than bacterial uptake rates (Fig. 3C), potentially due to either lower transporter affinities or lower expression, or both.

### NOSC as a predictor of substrate preferences and use efficiencies

Substrate oxidation state, NOSC, stands to directly impact preferences and SUE via thermodynamic constrains on internal metabolism or energetic payoffs (19). Thermodynamic regulation of metabolic rates is typically only considered when terminal electron acceptors, such as O_2_, are limiting, but there is growing support this phenomenon would also occur in carbon-limited, oxic environments (29). This regulation arises from the higher energy requirement (ΔG°*_Cox_*) of the oxidation half reaction as the substrate becomes more reduced (lower NOSC). The defined media used here can be inferred to be C-limited due to excess NH_4_^+^ supplied and since most amino acid LMWOS provide C:N at a ratio lower than biomass production demands.

Support for thermodynamic limitations from our substrate uptake observations are tenuous. We observed a negative correlation between NOSC and normalized substrate *t_50_*, indicating an earlier and more rapid uptake of oxidized (NOSC ≥ 0) organic acids, but only for the two bacterial species (Fig. 2A-B). The early utilization of oxidized substrates by bacterial species was further corroborated in the labeled experiments, where we observed the appearance of ^13^CO_2_ from LMWOSs with a higher NOSC (e.g., formate and acetate) before other labeled substrates with lower NOSC (Fig. 4D, 4E). While there can be a decoupling of substrate uptake and metabolic use in certain situations, this is a reasonable indicator of uptake. In instances where extracellular and respiration data were both present, *t_50_* values were similar or respiration data was slightly delayed. For the fungal species, there was no clear trend between NOSC and the midpoint of substrate depletion (*t_50_*) from unlabeled (Fig. 2C) or labeled experiments (Fig. 4F). It is unclear why there was not early uptake of more oxidized, organic acids by *P. spinulosum*, but this may reflect a metabolic strategy prioritizing sugar uptake (glucose, galactose, xylose). The potential trends we observed from unlabeled and labeled experiments for the two bacteria (Fig. 2A-B, Fig. 4D-E) provide some population-level support for whole community observations of faster uptake and mineralization of oxidized carboxylic acids (succinic acid, malic acid, formic acid) (15).

There were no significant trends supporting thermodynamic limitations on SUE (i.e., negative correlation between NOSC and SUE) (Fig. 2. 4G-I). For all isolates, formate had the lowest SUE (< 0.31), which is comparable to low anabolism/catabolism estimates in soil (15). SUE estimates of the two amino acids (glycine and valine) were higher than expected for the bacterial species under a hypothesized NOSC-SUE relationship (Fig. 4G-H) but similar in magnitude to previous observations (64). These high SUE values could be due to the demand for direct incorporation into proteins during rapid growth (65, 66). For the bacterial isolates, the disconnect between the overall relationship between substrate respiration dynamics and SUE (Fig. 4D, 4E) and the lack of a significant relationship between NOSC and SUE (Fig. 4G, 4H) may reflect a stronger control by metabolic entry points of the chosen LMWOS traced (39, 64, 66). The low SUE estimates for formate, for example, may be a result of the limited metabolic entry points for formate-derived carbon, which is assimilated by heterotrophs typically after conversion to CO_2_ by formate dehydrogenases (67, 68). Thus, formate utilization by these three isolates represents heterotrophic CO_2_ fixation via anaplerosis and, while still a significant route of C utilization (69), comprises a drastically different metabolic usage than the assimilation of the other labeled carbon substrates in organic form. While used previously in assessments of NOSC and SUE relationships (15), the distinct metabolic entry pathway for formate and inherently low incorporation of C via anaplerosis by heterotrophs (69) may be driving observed correlations between NOSC and SUE. Any future assessments of substrate NOSC should include alternative oxidized substrates (e.g., oxalate) that are known to be incorporated directly (70).

The lack of consensus across all three isolates implies NOSC is not a reliable parameter for predicting substrate utilization patterns or SUE. While predicted relationships between NOSC and LMWOS *t_50_* from bacterial observations agree with those produced in other stable isotope tracer experiments in soils (15), the population-level trends we observed were not strong (adj R^2^ < 0.12) and not present in data from the fungal isolate (Fig. 5). It remains unknown if the earlier use of oxidized organic acids than other substrates by these bacteria, whether a result of metabolism regulation or transporter affinity, would still be apparent with unequal substrate concentrations as is typical in soil solution (32). Previous attempts to connect the processing of LMWOS in soil to inherent chemical properties, such as C:N or molecular weight, have provided inconclusive results (71). Similarly, our results stress that a fuller understanding of the internal biochemical pathways, and not substrate NOSC alone, is necessary to predict LMWOSs cycling in soils (66).

**FIG 5.**
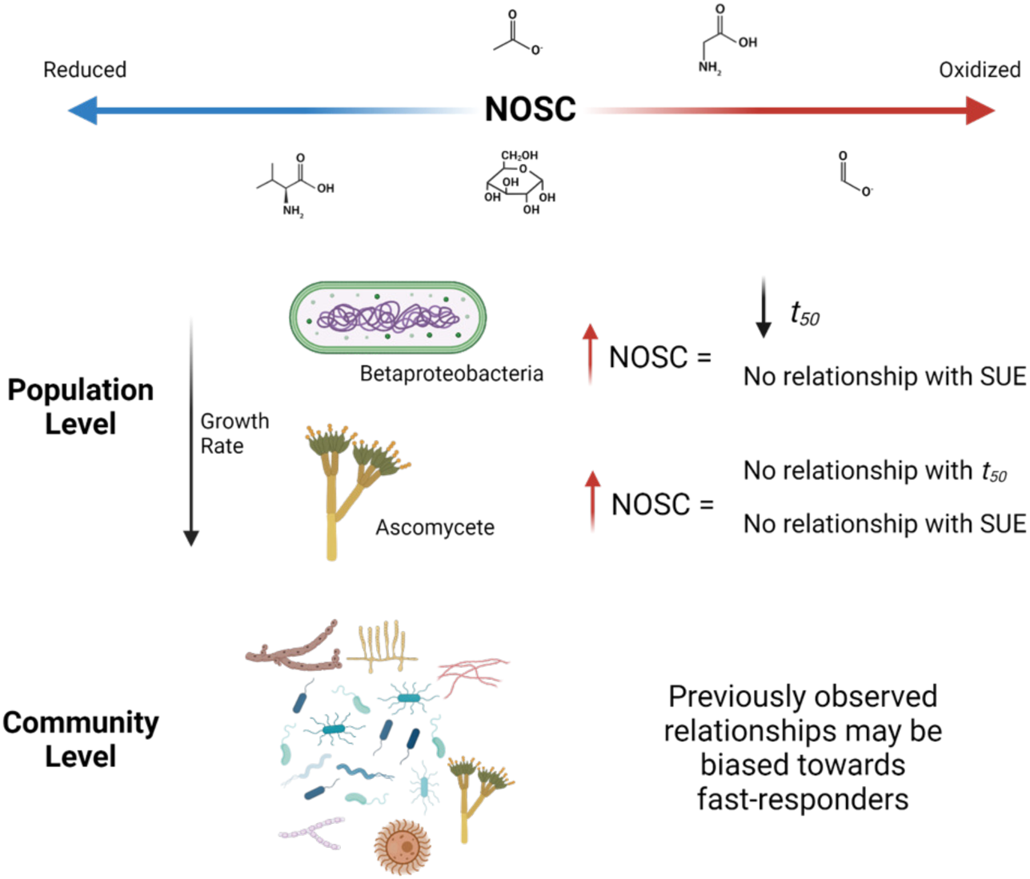
Summary chart outlining the results of this study and the possible implications at the community level. Created with BioRender.com.

Tracking the transformation of LMWOSs into soil carbon pools with longer turnover times requires knowledge of the principles governing microbial uptake and metabolic transformation. Many current conceptual frameworks for understanding these processes revolve around tradeoffs between microbial growth rate and CUE and the energy content of the LMWOS. Clustered substrate utilization was observed for all three microbial isolates cultured, indicating metabolic co-utilization of LMWOSs during growth. We only found potential trends (p < 0.05) supporting the preferential uptake of LMWOSs with higher NOSC for the two Betaproteobacteria, though variability was high and the overall effect of NOSC was small. We found no support for relationships between NOSC and SUE, highlighting the need to understand better the intracellular metabolism of LMWOS-carbon during growth. Despite community-level observations depicting a negative relationship between NOSC and half-life in solution as well as NOSC and SUE, our findings suggest that these observations may reflect the primary involvement of rapidly responding bacterial populations and that extrapolation of these results to other aerobic contexts should be approached with caution. Our ecophysiological approach, probing the carbon usage of diverse LMWOSs at realistic soil solution concentrations, presents a platform to test foundational relationships between LMWOS identity and substrate use across a range of microbial phylogenetic diversity and environmental contexts.

## Materials and Methods

### Microbial isolation and characterization

All strains were isolated from both field moist Oa and B horizons found under a hemlock-dominated stand in Arnot Forest, New York, USA. Three microbial strains were chosen from a library originally isolated using soil-extractable, solubilized organic matter (SESOM). All SESOM was derived from the Oa horizon. The extract was created using a modified water extraction (72), involving the suspension of 40 g of field-moist Oa horizon with 200 mL of 18.2 MΩ-cm water. Samples were shaken end-to-end for 1 hour at room temperature and then left to settle. After 24 hours, the solution was sequentially filtered through 1.6 μm GF/A, 0.45 μm polyethersulfone (PES), and then filter-sterilized using 0.2 μm PES filters. Enrichment steps were conducted using fresh soil shaken with DI water (1:10 ratio) for 1 hour and then left standing for 24 hours at room temperature. Enrichments were serially diluted using Winogradsky salts (73) and a 100 μL sample was then spread onto agar plates (15 g/L) created with SESOM as the sole C-source (denoted “N”) and with the addition of streptomycin and Rose Bengal (denoted “RB”). All plates were incubated in the dark at room temperature for 3-14 days. Colonies were chosen and re-streaked 3 times before growth was checked in SESOM liquid culture (2x dilution). Further details about the chemical characterization of SESOM and the growth of *Paraburkholderia solitsugae* in SESOM have previously been reported (32).

Isolates B3N, B1N, and Oa1RB were chosen based on growth rates in defined media and characterized using a combination of genomic approaches. Both bacterial species were identified using genomic DNA as discussed previously in Cyle et al., 2020. Briefly, genomic DNA was extracted from pelleted cells and submitted to the Cornell University Sequencing Facility for sequencing using three multiplexed runs of llumina MiSeq Nano (2 x 250 bp). Fungal identification was conducted using ITS1F (CTTGGTCATTTAGAGGAAGTAA) and ITS4 primers (TCCTCCGCTTATTGATATGC) (74). *Paraburkholderia solitsugae* was extensively characterized in prior works (32, 75). The other two isolates chosen are referred to by the species name with which they have the highest similarity (Table S1). Isolate B3N was found to be most related to *Ralstonia pickettii*, a rod-shaped Betaproteobacterium (76), while isolate Oa1RB was found to be most related to *Penicillium spinulosum*, an ascomycete (77).

### Media preparation and culturing conditions

All experiments were conducted in batch cultures using acid-washed and autoclaved 125-mL Erlenmeyer flasks using three biological replicates per treatment. Incubating flasks were maintained at room temperature on a shaker at 150 rpm. Defined media was prepared based on potential carbon substrates utilized by *P. solitsugae* during growth in SESOM (32). To assess the relative role of NOSC on utilization profiles and use efficiencies of each substrate, all carbon substrates were added to the media at equimolar concentration (25 μM each). Substrates included three sugars (glucose, galactose, xylose), 20 amino acids (alanine, arginine, asparagine, citrulline, cysteine, glutamate, glutamine, glycine, histidine, lysine, methionine, ornithine, isoleucine, leucine, lysine, phenylalanine, proline, serine, threonine, tryptophan, valine) and 10 organic acids (acetate, α-ketoglutarate, citrate, formate, gluconate, lactate, malate, oxalate, pyruvate, succinate). The defined medium was filter-sterilized and subsequently supplemented with 1.68 mM NH_4_Cl, 0.12 mM KH_2_PO_4_, 1x Wolfe’s vitamins and 1x Wolfe’s minerals solutions (78), and pH-adjusted to pH 4.5. The nutrient media was prepared in two different manners: (1) with all unlabeled substrates and (2) with a single substrate isotopically labeled with the remaining unlabeled substrates to allow the tracking of labeling into respired CO_2_ and into biomass.

Species were individually cultured in unlabeled defined media until exponential growth phase and then a sample of this culture was used to inoculate treatment flasks for experimental trials. For trials with bacterial species (*R. pickettii*, *P. solitsugae*), this involved overnight culturing and monitoring of growth using optical density at 595 nm (OD_595_) followed by inoculation of 50 mL treatment flasks with less than a 250 μL subsample (theoretical starting OD_595_ ∼ 0.0005). For trials with *P. spinulosum*, biomass measurements were made using destructive sampling of biological replicates of starter flasks (n = 3 per time point), filtration through 0.2 μm PES filters, and mass determination after drying at 55°C for 1 hour. At roughly the inflection point of the growth curve, a single flask was sonicated for 5 minutes and 100 μL volumes were used to initiate all treatments. All culturing work was conducted in a laminar flow hood using aseptic technique, sterile filter-pipette tips, and with negative controls (n = 3) to ensure sterility was maintained.

### Unlabeled growth trials. (I) Time-resolved metabolic footprinting sampling

For determining substrate utilization profiles, growth trials were conducted solely using the defined media with unlabeled substrates. A growth trial was initiated for each isolate as described above, with sufficient replicates to allow at least four destructive sampling time points across the organism’s growth curve. For bacterial species (*R. pickettii, P. solitsugae*), destructive sampling consisted of centrifugation of ∼40 mL of the culture at 10,000 x *g* for 10 minutes. Supernatant was decanted and filtered through 0.2 μm PES filters. For the fugal species (*P. spinulosum*), the entire culture of ∼50 mL was filtered through 0.2 μm PES filters. All samples were immediately allocated into 2-mL centrifuge tubes and stored frozen (−20°C) until exometabolomics analysis. All filtered samples were analyzed using a Shimadzu TOC-V_CPN_ for non-purgeable organic carbon (referred to as total organic carbon – TOC) and total nitrogen (TN) using a 2% acidification (0.2 M HCl) and 1:30 min sparge time using high temperature (720°C) catalytic (Pt) oxidation.

### (II)#Targeted analytes via LC-HRMS

Stored samples were thawed and prepared for immediate analysis using liquid chromatography coupled with high resolution mass spectrometry (LC-HRMS) as described in (32). Briefly, samples were analyzed using a Thermo Scientific Dionex Ultimate 3000 liquid chromatography system connected to a Q Exactive orbitrap mass spectrometer. A reversed-phase approach using a C18 column and negative electrospray ionization (79, 80) as well as a hydrophilic interaction approach using polarity switching (81) was used to quantitate substrate concentrations in the extracellular media. Quality control checks were run every 10 samples with a 30% standard deviation limit. All data was processed using an internally constructed template within Thermo Scientific Xcalibur 3.0 Quan browser using standards of all identified compounds run between 0-25 μM. Reliable data on three compounds not intentionally added to the minimal media (fumarate, cystine, and homoserine) have been included in all analyses.

### (III)#Targeted analytes via ^1^H NMR

Samples were analyzed using proton nuclear magnetic resonance spectrometry (^1^H NMR) to capture a select group of sugars and organic acids from the media that could not be quantified during some LC-HRMS analytical runs (glucose, galactose, xylose, acetate, formate, oxalate, valine). Methods used have been previously reported for extracted soil solutions (32, 72, 82). Briefly, 35-40 mL samples of frozen extracellular media were concentrated by lyophilizing the sample and reconstituting to a smaller, final volume of 500 μL. Reconstitution involved 300 μL of 18.2 MΩ-cm water and buffered to pH of 7.0 using an addition of 200 μL of sodium hydrogen phosphate (0.1 mM, pH 7.0) made with 25% D_2_O (vol/vol) to supply a lock signal and containing 1 mM sodium 3-trimethylsilyl-[2,2,3,3,-D_4_]-1 propionic acid (TMSP) to provide spectral referencing at a final concentration of 0.4 mM. All spectra were collected at 500 MHz at room temperature on a Bruker AV 500 operated using Bruker TopSpin 3.5.7 using a 10% D_2_O and water peak suppression program (noesygppr1d) with 32 scans/sample and a 5-s relaxation delay for a total of 256 transients. Previously described spectral processing methods and integral regions (32) were used on all samples. Areas under the curve were normalized to initial media values for each experiment and used for depletion model fitting. No reliable sugar data was able to be collected, though in some cases valine, acetate, and formate were able to be modeled and are included in analyses.

### (IV)#Curve fits for microbial growth and substrate depletion analysis

Microbial growth was modeled using R 3.6.0 (83) using the nls.multstart package (84). Microbial growth was modeled in terms of biomass (mg L^−1^) to ensure equivalent comparison of growth rates between bacterial isolates, which were originally monitored in terms of OD_595_, and the fungal isolate, which was originally monitored in terms of biomass. Conversion from OD_595_ to mg L^−1^ for the bacterial species was conducted using a biomass conversion factor (*k* – mg L^−1^, Table 1), determined on and OD unit basis using biomass data collected at stationary phase during the labeled growth trials. Growth was modeled using a reparametrized Gompertz equation (Eq. 1) (85) on untransformed biomass data to allow extraction of parameters with biological meaning and for visualization purposes,

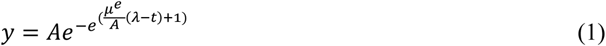

where *y* is average biomass data (mg L^−1^), *A* is the stationary phase asymptote of the growth curve (mg L^−1^), *t* is time (h), μ is the specific growth rate (h^−1^), λ is the lag time (h^−1^). A secondary fitting approach was employed using the growthrates package (86) to produce normalized estimates of specific growth rate (μ, hr^−1^) from average biomass data, readily allowing comparisons with literature values while avoiding biases from estimates of biomass from initial cell density (N_0_) (87).

Substrate depletion was analyzed based on pattern clustering, nonlinear modeling, and calculations of maximum depletion rate. Clusters of substrate depletion were identified for each isolate using a k-means approach specifically suited for comparing trajectories. Substrate depletion observations, normalized to measured initial media concentrations, were analyzed using the kml package (88) in R 3.6.0 (83). Clusters were iteratively analyzed ensuring clusters of sufficient size (10%) and maximizing the Calinski-Harabanz index (89). Model fits of substrate depletion data and extracellular TOC were created for individual biological replicates using the nls.multstart package (84) as described previously (32, 40, 90). A nonlinear modeling approach (Eq. 2) allowed the fitting of a 4-point sigmoidal curve using the following equation:

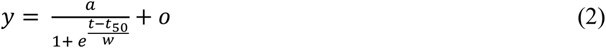

where *y* represented either concentration data (μM) or in some cases was normalized relative to initial conditions (0-1) and *t* represents time (h). The four parameters produced during the fitting procedure relate to the amplitude of substrate depletion (*a*, μM or unitless), the midpoint of depletion (*t_50_*, h), the width of the concentration decrease (*w*, h), and the offset or predicted final value of substrate remaining in extracellular media (*o*, μM or unitless). For visualization and comparison purposes, the midpoint of depletion (*t_50_*) has been chosen as the point to assess substrate uptake ordering. The width parameter has been modified in all figures to show a wide usage window towards capturing substrate use overlap. The usage window extends from 10% of total substrate utilization or where y = [(*a*-*o*) x 0.9] as well as the time at which 90% of total substrate utilization has occurred or where y = [(*a*-*o*) x 0.1]. In some cases, no fit was applied due to insufficient data or indications that a sigmoidal fit was not the appropriate model.

Maximum depletion rate was determined by taking the differential of the modeled substrate depletion curve and calculating the rate at the inflection point (*t_50_*) (40) and expressing that value as a rate normalized in terms of cell dry weight (μmol h^−1^ mg_CDW_). In some cases, the midpoint of depletion, *t_50_*, has been normalized to the stationary phase sampling time (0-1) for easier comparison across isolate growth curves. All two-group comparisons between compound classes were conducted using a Welch’s t-test and the relationship between NOSC and normalized *t_50_* was assessed using linear regression.

### Labeled growth trials. (I) Respiration sampling during growth

Growth trials were conducted for each isolate using labeled substrates (> 98% ^13^C, 99% ^15^N, Cambridge Isotope Laboratories, Inc.) to track C into ^13^C labeled microbial biomass as well as respired ^13^CO_2_. Five labeled treatments were chosen to span low molecular weight compound classes (sugar, organic acid, amino acid) as well as nominal oxidation states of carbon (−0.8 – 2). Substrates included a sugar (glucose, NOSC = 0), two organic acids (acetate, NOSC = 0 and formate, NOSC = 2), and two amino acids (glycine, NOSC = 1 and valine, NOSC = −0.8). Each labeled treatment contained the labeled substrate as well as the remaining unlabeled 33 substrates.

Culturing was conducted as described previously except rubber septa were fitted after inoculation to seal off the headspace of the flask. At intervals during growth, 250 μL headspace gas samples were taken using a 500 μL gastight syringe. Gas samples were then injected into pre-evacuated, and helium (He) filled 2 mL glass crimp vials sealed with a PTFE/butyl septum. The quantity of ^12^CO_2_ (m/z 44) and ^13^CO_2_ (m/z 45) were measured using gas chromatography-mass spectrometry using a Shimadzu GCMS-QP2010S equipped with a Carboxen 1010 PLOT column and ultra-high purity He (Airgas, Inc.) as the carrier gas using previously described protocols (18, 91). Samples were run within 24 hours and each sample run was accompanied by standards ranging from 0-17,700 ppm created using CO_2_ (Airgas, Inc.). No purging was conducted in between sampling points, so all data represents cumulative buildup of CO_2_ in the headspace. All values are displayed as CO_2_-C (% of addition). This was calculated by first subtracting natural abundance ^13^CO_2_ determined using the unlabeled control treatments. This enriched ^13^CO_2_ value was converted to mg ^13^C and divided by the amount of labeled substrate supplied in the treatment. Cumulative CO_2_ curves were modeled using a modified version of equation 2 with the numerator altered to [-(x – *t_50_*)] to invert the 4-point sigmoidal fit for the hypothetical curve of mirrored CO_2_ release. The offset parameter (*o*) was also set to 0 for this scenario.

### (II)#Biomass harvest and isotopic measurements

At the beginning of stationary phase, isolate biomass was destructively harvested from all culture flasks. Destructive sampling was conducted as described for unlabeled growth trials. All biomass, whether separated using a filter or centrifugation, was washed using 5 mL of C-free 1x Wolfe’s minerals solutions. A portion of separated biomass was then submitted to the Cornell Stable Isotope Laboratory for combustion analysis using a Thermo Delta V isotope ratio mass spectrometer (IRMS) interfaced to a NC2500 elemental analyzer.

### (III)#Calculations and data visualization

All post-processing was conducted in R 3.6.0 (83) using the tidyverse package (92), the RColorBrewer package (93), and the cowplot package (94). The SUE (Eq. 3) and CUE (Eq. 4) values were both estimated by combining the isotopic data for CO_2_ and biomass as evaluated at stationary phase. The SUE was calculated as:

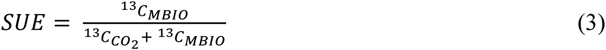

where ^13^C_CO2_ (mg) and ^13^C_MBIO_ (mg) was determined for each replicate after subtraction of the average natural abundance ^13^C in both pools using atom percent ^13^C values from unlabeled treatments. The CUE was calculated similarly but using the total values of C.

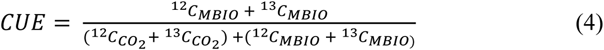

Statistical comparisons between treatments for dependent variables (*t_50_*, SUE) were analyzed first using ANOVA and then subsequent multiple comparison tests were conducted using Tukey’s HSD. Correlations were assessed using linear regression.

### Data Availability

The genome assembly for strain B1N (*Paraburkholderia solitsugae*) and strain B3N (*Ralstonia pickettii*) can be accessed via the NCBI portal using the BioProject accession numbers PRJNA590275. The ITS sequence for strain Oa1RB (*Penicillium spinulosum*) can be accessed via the NCBI GenBank portal using accession numbers MZ375756. Metabolomics data have been deposited to the EMBL-EBI MetaboLights database (DOI: 10.1093/nar/gkz1019, PMID:31691833) with the identifier MTBLS3558 (95).

## Acknowledgements

Graduate financial support for K.T.C. was provided by the College of Agriculture and Life Sciences at Cornell University. Partial graduate financial support and funding was provided by the Cornell University Program in Cross-Scale Biogeochemistry and Climate, which is supported by NSF-IGERT and the Atkinson Center for a Sustainable Future. Postdoctoral support for A.R.K. was provided by a National Science Foundation CAREER Grant (award # 1653092) awarded to L.A. This work was also supported by the AFRI Education and Workforce Development Program, grant no. 2019-67011-29513, from the U.S. Department of Agriculture, National Institute of Food and Agriculture. We thank Roland Wilhelm, Dan Buckley, and Martínez Research Group members for helpful revisions to the manuscript. Sequencing was performed by the Biotechnology Resource Center (BRC) Genomics Facility at Cornell University (http://www.biotech.cornell.edu/brc/genomics-facility). ^1^H NMR experiments were performed at the Cornell University NMR facility in the Department of Chemistry and Chemical Biology.

